# The orthobunyavirus Gc glycoprotein head and stalk drives an infectious virion assembly pathway that is specific for the insect host

**DOI:** 10.1101/2025.11.18.689178

**Authors:** Amelia B. Shaw, Hiu Nam Tse, Molly V. Durawa, Owen Byford, Hayley Pearson, Kenneth Stapleford, Juan Fontana, John N. Barr

## Abstract

The *Orthobunyavirus* genus of arthropod-borne segmented RNA viruses comprises several important pathogens including the human-infecting Oropouche virus and animal-infecting Schmallenberg virus (SBV). The prototypical Bunyamwera orthobunyavirus (BUNV) possesses envelope-embedded glycoprotein spikes, with Gc head and stalk domains forming distinctive tripods covering the envelope-proximal Gn. Spike ectodomains mediate virus entry, while endodomains interact with nucleoprotein (NP) enwrapped genome segments to mediate virion assembly. Interestingly, BUNV Gc head/stalk domains are redundant for virus growth in mammalian cells, consistent with isolations of SBV from animals bearing head/stalk deletions. However, orthobunyavirus isolations from arthropods in nature appear to maintain these domains strictly.

To investigate this discrepancy, we compared the multiplication characteristics of wildtype BUNV (BUNV-WT) with a Gc head/stalk deleted BUNV (BUNV-Δ7). In mammalian cells BUNV-WT and BUNV-Δ7 grew to equivalent titres, whereas BUNV-Δ7 titres from insect cells were 1000-fold lower than BUNV-WT and strikingly produced no virions following blood meal infection of *Aedes* mosquitoes.

To understand this insect-specific restriction in virion production, we showed the intracellular abundance of BUNV-WT and BUNV-Δ7 Gc and NP components were equivalent, suggesting Δ7-Gc was assembly-deficient. To explore this, we investigated Gc and Δ7-Gc interactions during BUNV-WT and BUNV-Δ7 infections of both insect and mammalian cells by co-immunoprecipitation and multiplex mass spectrometry, revealing Δ7-Gc exhibited markedly reduced NP interactions in insect cells. We propose Gc recruits genome segments during virion assembly and that the complete Gc head/stalk assembly is maintained in nature due to its essential role in the insect host.

**IMPORTANCE:** Orthobunyaviruses are arthropod-borne viruses that cause severe disease in humans and animals, including congenital malformations and abortions. Orthobunyavirus Gn and Gc glycoproteins form spikes, which mediate entry and genome recruitment during assembly. In nature, OBVs bearing large deletions within Gc head/stalk domains have been isolated from animals, yet the head/stalk domains appear to be strictly maintained within insects. To investigate this discrepancy, we compared the multiplication of Bunyamwera orthobunyavirus (BUNV-WT) with head/stalk-deleted variant (BUNV-Δ7). In mammalian cells both viruses reached similar titres, but strikingly BUNV-Δ7 failed to produce virions in both insect cells and *Aedes* mosquitoes. We showed this was because BUNV-Δ7 failed to assemble new virions, revealing its head/stalk-deleted Gc was deficient in interactions with genome components. We propose that Gc drives species-specific interactions with genome segments during virion assembly, explaining why Gc head/stalk domains are conserved in nature due to their essential role in the insect host.

## INTRODUCTION

The *Orthobunyavirus* genus within the *Bunyaviricetes* class is a large group of arthropod-borne segmented negative-sense RNA viruses that inflict a significant disease burden in mammals. Bunyamwera virus (BUNV) is the prototype of the orthobunyavirus (OBV) group and causes febrile illness in humans, although the BUNV reassortant Ngari virus has been associated with fatal outbreaks of haemorrhagic fevers (Heitmann et al., 2021). The OBV group also includes Oropouche virus (OROV), capable of placental infection with subsequent foetal injury including microcephaly, and La Crosse virus, responsible for widespread outbreaks of encephalitis. Several OBVs also have a significant impact on animal health, including Aino virus (AINOV), Akabane virus (AKAV) and Schmallenberg virus (SBV), which are capable of vertical transmission from mother to neonate, resulting in severe congenital malformations and foetal abortion. All OBVs are maintained in nature by a vertebrate-arthropod transmission cycle, in which persistently infected arthropods act as vectors, transmitting the virus to vertebrates during a blood meal.

The OBV genome comprises three segments; the S segment encodes the nucleocapsid protein (NP) and the non-structural S protein (NSs). The M segment encodes a glycoprotein precursor (GPC) which is co-translationally cleaved to yield the glycoproteins Gn and Gc, and the non-structural M protein (NSm). Finally, the L segment encodes the large protein (L) which represents the virus-encoded component of the RNA-dependent RNA polymerase (RdRp). All three RNA segments associate with the RdRp and are protectively encapsidated with NP forming helical and pseudo-circular structures known as ribonucleoprotein (RNP) complexes (Hopkins et al., 2022).

OBVs, like other bunyaviruses, enter mammalian cells using their glycoprotein spikes, which protrude from the viral envelope. The spikes interact with various cellular receptors, such as lipoprotein-related protein 1 (LRP-1) recently identified as the receptor for OROV (Schwarz et al., 2022). Following internalization into endosomes, the spikes also direct fusion of the viral and host membranes resulting in release of RNP segments into the cytosol where transcription and subsequently replication occur (Hover et al., 2018, 2023; Punch et al., 2018). In mammalian cells, assembly of new OBV particles occurs within a re-organised Golgi, where newly made RNPs associate with the cytoplasmic tails of Gn/Gc glycoproteins (Shi et al., 2007; Strandin et al., 2013). A similar mechanism has been observed for other bunyaviruses (Överby et al., 2007; Piper et al., 2011). Nascent OBV virions form and accumulate within the Golgi lumen to subsequently bud into exocytic vesicles and mature during trafficking to the plasma membrane for release (Fontana et al., 2008; Novoa et al., 2005; Salanueva et al., 2003). The OBV infection cycle within insect cells is less well understood, although BUNV does not cause a similar ultrastructural reorganisation or swelling of the Golgi, with newly generated BUNV virions proposed to be released from insect cells immediately after formation without accumulation, facilitating long-term persistent infections (Elliott & Wilkie, 1986; López-Montero & Risco, 2011).

The OBV Gc protein is sub-divided into three structural domains: the head, stalk and floor. The ultrastructural organisation of the BUNV glycoproteins on the viral surface has been characterised, displaying a tripodal organisation. Here, three Gc head domains form the apex of the tripod, connected to the base of the tripod through the stalk domain (Bowden et al., 2013; Hellert et al., 2019; Hover et al., 2023). The tripod base is composed of three Gc floor domains, which contain the class II fusion domain, alongside three copies of Gn, forming a trimer of heterodimers, which stabilises the tripodal structure and shields the fusion domain (Hover et al., 2023). Both Gc and Gn possess C-terminal cytoplasmic tails that protrude inside the virion, and in the absence of a canonical viral matrix protein, have been proposed to interact with the RNPs, driving virion assembly and budding (Shi et al., 2007).

The Gc head region is highly variable and several viable OBV variants with mutations and deletions in the Gc head region have been identified. Headless variants were first described in 1981 during passage of Maguari virus (MAGV) in mammalian cells in the presence of the mutagen 5-fluorouracil (Iroegbu & Pringle, 1981; Pollitt et al., 2006). Mutations in the SBV Gc head domain were also identified following serial passage in mammalian IFN-incompetent cell lines (Varela et al., 2016). For BUNV, a systematic deletion analysis of Gc revealed the head/stalk domains to be largely dispensable for multiplication in mammalian cells (Shi et al., 2009) with one variant, termed delta-7 (Δ7), exhibiting similar growth kinetics to wild-type BUNV despite removal of the entire head domain and most of the stalk region (Shi et al., 2009).

Interestingly, viruses bearing extensive Gc mutations have also been isolated in nature. For SBV, isolates extracted from ruminants revealed a mutational hotspot in the Gc head domain (Fischer et al., 2013) and later studies identified multiple SBV mutants bearing deletions of up to 200 amino acids, with some encompassing portions of both Gc head and stalk domains (Coupeau et al., 2013; Fischer et al., 2013; Wernike et al., 2021). Interestingly, no isolations of Gc-deleted OBVs from arthropods have been reported, suggesting the functional requirements of Gc may be different across the two hosts that make up the OBV sylvatic multiplication cycle.

Here, we wanted to further investigate the role of the OBV Gc head and stalk domains, to better understand why they are apparently dispensable for multiplication in the mammalian host, yet are maintained in nature. Using BUNV, we examined the growth of both rBUNV-WT and rBUNV-Δ7 in mammalian-derived A549 and BHK cells, alongside arthropod-derived C6/36 cells. While structural proteins Gc and NP from both viruses accumulated with comparable abundance in all three cell lines, in insect cells, we observed redistribution of Δ7-Gc to the plasma membrane alongside a dramatic drop in released rBUNV-Δ7 titres by over 1000-fold. These observations suggested rBUNV-Δ7 was deficient in virion assembly, recapitulated by the failure of rBUNV-Δ7 to multiply in mosquitoes. To investigate the basis of this deficit we used co-immunoprecipitation paired with tandem mass tag (TMT) mass spectrometry to examine Gc interactions within infected cells. This revealed a significant reduction in the ability of the head/stalk truncated Gc of rBUNV-Δ7 to interact with NP in insect cells, which was not seen in mammalian cells. This suggests Gc potentially plays a role in RNP recruitment and its deletion blocks virion assembly within the insect host. These findings provide a plausible explanation for why Gc-truncated OBVs can be isolated from mammalian hosts, but not from arthropods.

## RESULTS

### rBUNV-Δ7 exhibits no fitness loss in mammalian cells but is blocked in virus production in insect cells

To better understand the role of the Gc head/stalk domain in OBV growth, we compared the replication characteristics of wildtype BUNV and the variant rBUNV-Δ7 missing the Gc head/stalk region. To achieve this, we rescued recombinant wildtype BUNV (rBUNV-WT) and the variant rBUNV-Δ7, as previously described (Shi et al., 2009) (Supplemental Fig 1A), with successful rescue confirmed by western blot analysis of both transfected and infected cell lysates (Supplemental Fig 1B). To reflect the two-host enzootic multiplication cycle of OBVs, we compared rBUNV-WT and rBUNV-Δ7 activities in mammalian-derived A549 and BHK cells as well as mosquito-derived C6/36 cells.

To assess their multiplication characteristics, we performed a time-course experiment in which A549, BHK and C6/36 cells were infected with either rBUNV-WT or rBUNV-Δ7 at an MOI of 5, with both lysates and supernatants harvested at 6-, 12- and 24-hours post infection (hpi) (Fig 1). Lysates were probed for expression of NP using western blot analysis (Fig 1A-B) with subsequent densitometry revealing similar NP expression from both rBUNV-WT or rBUNV-Δ7 at all time points measured in A549 and BHK cells. In C6/36 cells, rBUNV-Δ7 NP appeared slightly earlier (Fig 1A-B), but this was not statistically significant, and overall NP expression of the two viruses at 24 hpi was indistinguishable. This suggested rBUNV-WT and rBUNV-Δ7 were able to enter and perform gene expression all three cell lines in an equivalent manner.

**FIG 1.**
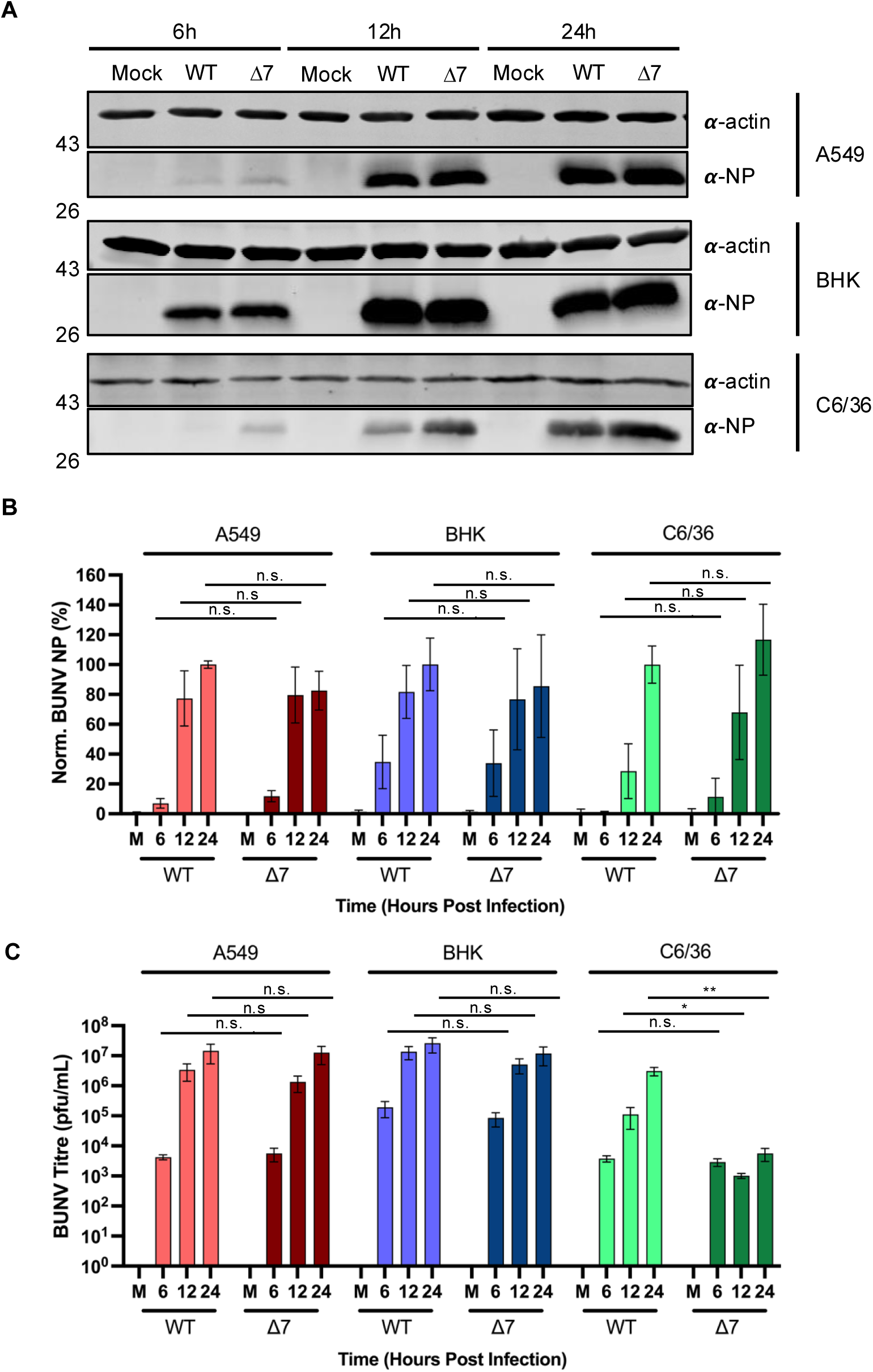
Comparison of growth kinetics between wildtype BUNV and Δ7 BUNV in multiple cell lines. A549 cells (mammalian), BHK cells (mammalian) and C6/36 cells (insect) were either mock-infected or infected with rBUNV-WT (WT) or mutant rBUNV-Δ7 (Δ7) at an MOI of 5 and supernatants and cell lysates were collected at either 6, 12 or 24 hours post infection (n=3). (A) Cell lysates were subject to western blot analysis and probed for expression of NP and actin, as a loading control. (B) Densitometry analysis performed on (A) as expression of NP relative to actin. The expression of BUNV NP was normalised to WT at 24 hours post infection for each cell line. Results were analysed by Student’s t-test (unpaired) whereby n.s. = p>0.05, comparing WT to Δ7 at each timepoint. (C) Titration of the supernatants was performed by plaque assay. Log_10_-transformed samples were analysed by Student’s t-test (unpaired) whereby n.s. = p>0.05, * = p<0.05; ** = p<0.01; *** = p<0.001, comparing WT to Δ7 at each timepoint.

Next, we assessed the titre of infectious rBUNV-WT or rBUNV-Δ7 released from the three cell lines (Fig 1C) in harvested supernatants. The same three timepoints (6, 12 and 24 hpi) as for lysate harvesting were used, with 6 hpi reflecting the baseline titre when no new virus was generated, 12 hpi reflecting virus released after just one round of infection, and 24 hpi reflecting virus released from multiple rounds of infection. Supernatant titre analysis by plaque assay showed release of infectious rBUNV-WT and rBUNV-Δ7 from mammalian A549 and BHK cells similarly increased over all three timepoints, reaching approximate titres of 10^7^ pfu/mL at 24 hpi. Interestingly, while this was comparable to the release of infectious rBUNV-WT from C6/36 cells, the titre of infectious rBUNV-Δ7 from C6/36 cell supernatants was reduced by around 1000-fold and in fact did not increase beyond the baseline titre (6 hpi) suggesting no new virus was released. This suggests that the head/stalk domain is essential for release of infectious BUNV from insect cells, but dispensable in mammalian cells.

### rBUNV-Δ7 cannot multiply in Aedes mosquitoes

To assess whether the assembly block in rBUNV-Δ7 infection was specific to insect cells in culture, we sought to test the multiplication characteristics of rBUNV-Δ7 in female *Aedes aegypti* mosquitoes. Briefly, over 20 mosquitoes were fed blood-meals containing either rBUNV-WT or rBUNV-Δ7 at 1 x 10^7^ pfu/mL, with day 0 samples collected immediately after feeding to confirm successful infection (Fig 2A; Day 0). At 7 days post infection (dpi), mosquitoes were collected and titres were assessed (Fig 2A; Day 7) from the bodies, to determine infection progress, and from the legs and wings, to determine effective dissemination. Of all the mosquitoes fed with a blood-meal containing rBUNV-WT, 10/21 mosquitoes were successfully infected with rBUNV-WT and in 9 of those, BUNV had successfully disseminated to the legs and wings. Of the mosquitoes fed with a blood-meal containing rBUNV-Δ7, 0/25 mosquitoes were infected with rBUNV-Δ7 nor was there any dissemination to the legs and wings. This was corroborated by western blot analysis of mosquitoes fed with either rBUNV-WT (10 mosquitoes) or rBUNV-Δ7 (10 mosquitoes), which showed 3/10 mosquitoes infected with rBUNV-WT expressed BUNV NP, whereas there was no NP expression in any of the mosquitoes infected with rBUNV-Δ7 (Fig 2B). This confirmed that the Gc head was critical to successfully establish infection in mosquitoes, corroborating our insect cell culture data.

**FIG 2.**
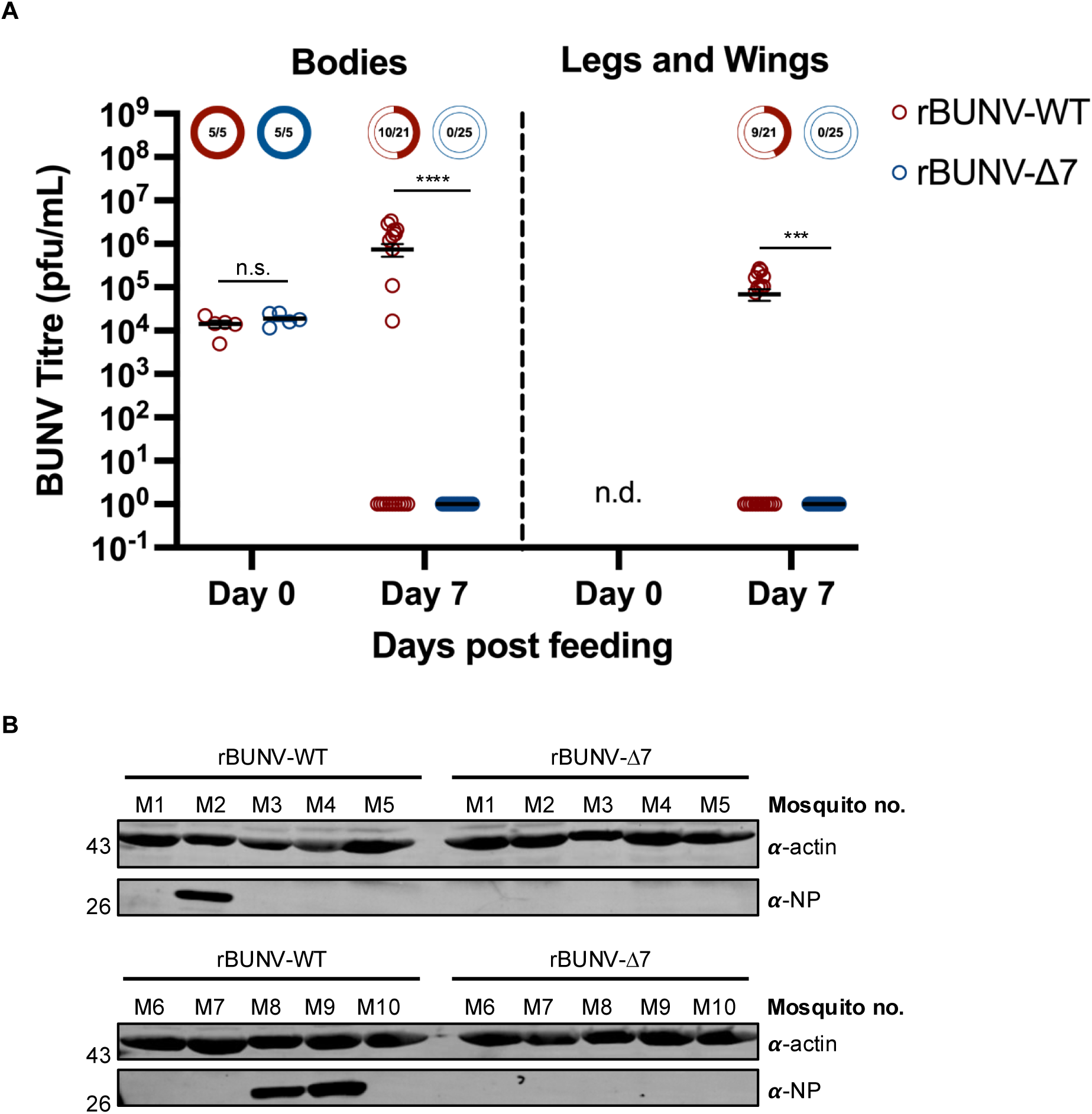
Comparison of infection of *Aedes* mosquitoes by wildtype BUNV and BUNV-Δ7. *Aedes* mosquitoes were fed a blood meal containing 1×10^7^ pfu/mL of rBUNV-WT or rBUNV-Δ7. (A) Bodies or legs and wings were harvested either immediately (Day 0) or 7 days later (Day 7) and titre was calculated by plaque assay. Titres are displayed as pfu/mL and individual samples have been shown (n=5 for both viruses at Day 0; n=21 for rBUNV-WT and n=25 for rBUNV-Δ7 at Day 7). The bar represents the average value, with standard error of the mean. Titres were analysed by Mann Whitney test (unpaired) whereby n.s. = p>0.05, *** = p<0.001, **** = p<0.0001, comparing WT to Δ7. (B) Mosquito lysates (10 mosquitoes for each virus) were subject to western blot analysis and probed for expression of NP and actin, as a loading control.

### rBUNV-Δ7 does not accumulate within insect cells or synthesise non-infectious virus-like particles

The lack of infectious rBUNV-Δ7 released into C6/36 cell supernatants could have been due to a defect in virion assembly, or a defect in virion release. To distinguish between these possibilities, we next assessed whether infectious virions were trapped inside cells. A549 and C6/36 cells were infected with either rBUNV-WT or rBUNV-Δ7 at an MOI of 5, after which titres of both supernatant-associated extracellular virus and cell lysate-associated intracellular virus were assessed by plaque assay at 24 hpi. The titres of extracellular rBUNV-WT released from A549 and C6/36 cells were equivalent (Fig 3A and B, extracellular), and consistent with previous 24 hpi supernatant titres, described above (Fig 1), as was the statistically significant reduction of titre of extracellular infectious rBUNV-Δ7 in C6/36 cells (Fig 3B; extracellular). Critically, the titre of C6/36 intracellular infectious rBUNV-Δ7 was statistically indistinguishable from that of rBUNV-WT (Fig 3B; intracellular), suggesting the dramatic reduction in rBUNV-Δ7 supernatant titres was not due to an intracellular accumulation of assembled virions. In A549 cells, there was no significant difference between titres of intracellular or extracellular infectious rBUNV-Δ7 and rBUNV-WT (Fig 3A).

**FIG 3.**
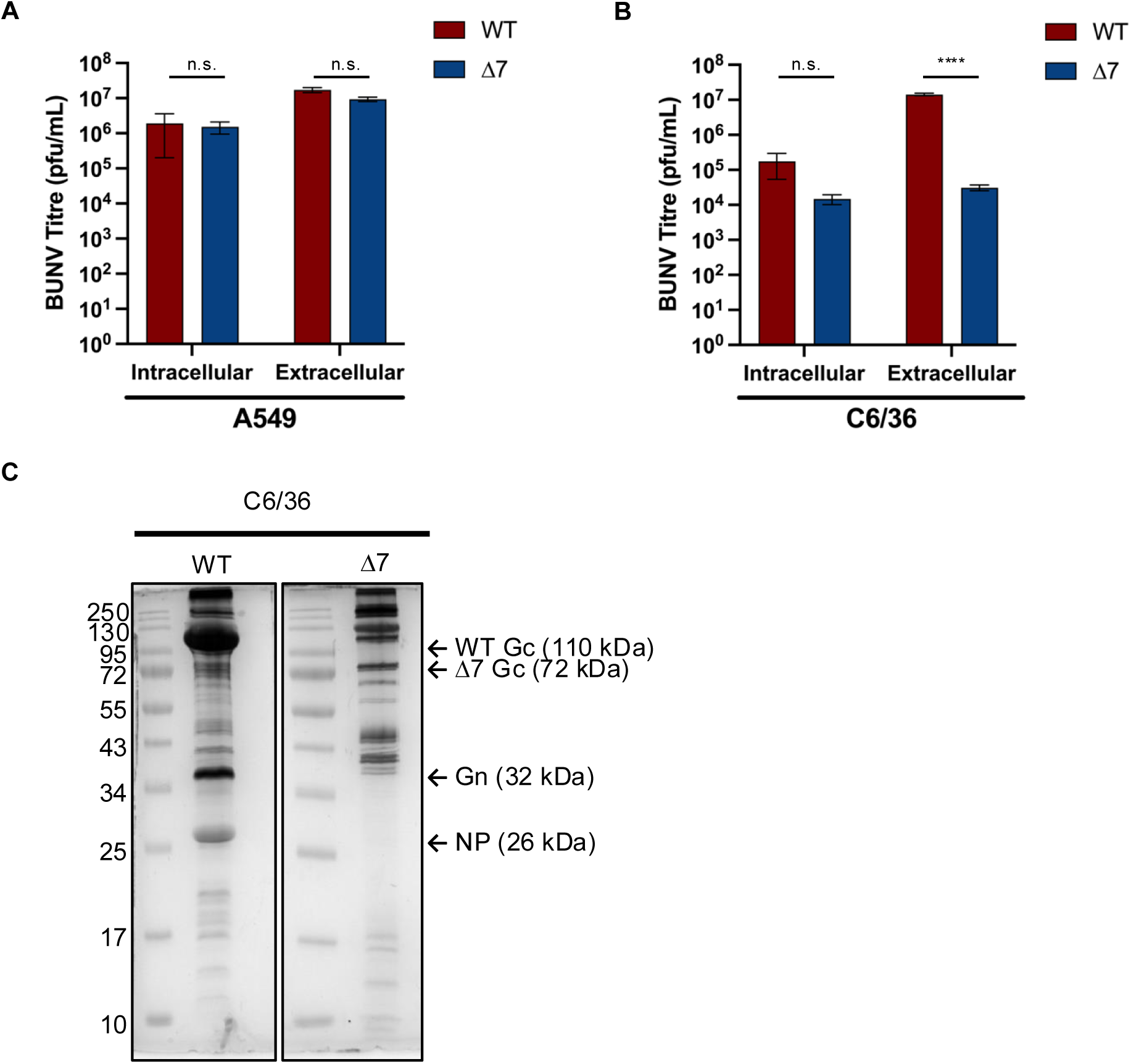
Comparison of intracellular and extracellular virus production of wildtype BUNV and Δ7 BUNV in A549 and C6/36 cells. A549 cells (A) or C6/36 cells (B) were infected with rBUNV-WT (WT; red) or mutant rBUNV-Δ7 (Δ7; blue) at an MOI of 5 and supernatants and cells were collected at 24 hours post infection (n=3). Cells were then freeze-thawed three times in 1x TNE buffer to disrupt the plasma membrane and release any assembled virions that were present within the cell (intracellular). Intracellular and extracellular (supernatant samples) were then titred by plaque assay. Log_10_-transformed samples were analysed by Student’s t-test (unpaired) whereby n.s. = p>0.05, **** = p<0.0001, comparing WT to Δ7 for each sample. (C) Extracellular supernatant from C6/36 cells infected with rBUNV-WT or rBUNV-Δ7 was purified by ultracentrifugation. The resuspended viral pellet was subject to SDS-PAGE and silver stain analysis. The protein ladder sizes are indicated (kDa) as well as the predicted bands for viral structural proteins WT-Gc, Δ7-Gc, Gn and NP (n=1).

As an orthogonal approach to detect accumulated intracellular virions, we directly observed sections of resin-embedded infected C6/36 cells by electron microscopy (Supplemental Fig 2). However, we were unable to detect any intracellular virions in C6/36 cells infected with either rBUNV-WT (López-Montero & Risco, 2011); Supplemental Fig 2B) or rBUNV-Δ7 (Supplemental Fig 2C).

Finally, to determine whether non-infectious rBUNV-Δ7 virus-like particles (VLPs) were released, we sought to examine presence of VLPs in infected C6/36 cell supernatant. Supernatant was collected from C6/36 cells infected with either rBUNV-WT or rBUNV-Δ7 and concentrated by ultracentrifugation through a sucrose cushion. The purified supernatant was examined by silver stain analysis (Fig 3C), which showed bands corresponding to the viral structural proteins Gc, Gn and NP for rBUNV-WT. However, the corresponding bands for rBUNV-Δ7 were much fainter, suggesting that VLPs are not being assembled in insect cells for this virus. The supernatant was also subject to cryo-electron microscopy, which showed a high number of rBUNV-WT virus particles but no similar virus particles from rBUNV-Δ7-infected C6/36 cells (data not shown), consistent with a lack of virus particle production from rBUNV-Δ7-infected C6/36 cells.

Taken together, our findings are consistent with a scenario in which lack of released infectious rBUNV-Δ7 within C6/36 cells results from a deficiency in virion assembly.

### Differences between insect and mammalian cells do not account for the reduction in rBUNV-Δ7 titres released from insect cells

To understand the molecular mechanism responsible for the lack of assembly and release of rBUNV-Δ7 from insect cells, we considered differences between the mammalian and insect cell lines, including lower culture temperature and lower cholesterol levels in the cellular membrane. To investigate whether reducing the culture temperature affected growth of rBUNV-Δ7, A549 and BHK cells were infected with either rBUNV-WT or rBUNV-Δ7 at an MOI of 5 and incubated at 30 °C, instead of 37 °C. At 24 hpi, supernatant titres were measured, which showed both rBUNV-WT and rBUNV-Δ7 reached equivalent high 10^6^ pfu/mL titres (Supplemental Fig 3A). This suggests that reducing the culture temperature did not account for the reduction of rBUNV-Δ7 titre in insect cells.

Unlike mammals, insects do not synthesise cholesterol, instead relying on dietary sources (Krebs & Lan, 2003). Given cholesterol abundance has been shown to regulate BUNV entry (Charlton et al., 2019), we hypothesised that differences in cholesterol incorporation in the cellular membranes of the mammalian and insect cell lines could account for the block in infectious rBUNV-Δ7 release. In an attempt to recover rBUNV-Δ7 viability, we supplemented C6/36 insect cell cultures with cholesterol during infection using an established cholesterol supplementation protocol (Thannickal et al., 2023). To do this, C6/36 cells were pre-treated with either 0.05 mg/mL or 0.1 mg/mL cholesterol, alongside methyl-β-cyclodextrin (MβCD; 85 µM) as a vehicle control, before infecting with rBUNV-WT or rBUNV-Δ7 at MOI 5 and harvesting supernatant at 24 hpi (Supplemental Fig 3B). Titration of the supernatants showed that supplementing C6/36 cells with cholesterol did not affect rBUNV-WT titre, as expected, nor did it recover rBUNV-Δ7 titre. This suggests that the absence of cholesterol in insect cells is not the reason for the reduction in released rBUNV-Δ7 titres.

### Glycosylation and trimerisation sites absent in rBUNV-Δ7 are not the reason behind the reduced rBUNV-Δ7 titres from insect cells

WT-Gc and Δ7-Gc sequences differ in their glycosylation potential due to the loss of residue N624 as a consequence of the head/stalk deletion. To investigate whether glycosylation differences could account for reduced rBUNV-Δ7 supernatant titres, we removed this glycosylation site from rBUNV-WT by the substitution N624Q to generate mutant rBUNV-N624Q. We hypothesized that if this residue played a critical role in infectious rBUNV-Δ7 production, then titres of rBUNV-N624Q would also be low. A549 cells or C6/36 cells were infected with rBUNV-WT, rBUNV-Δ7 or rBUNV-N624Q at MOI of 1 and incubated for 24 hours. Titration of the supernatant showed that rBUNV-WT and rBUNV-Δ7 were again similar in A549 cells and the rBUNV-Δ7 titre was reduced in C6/36 cells (Supplemental Fig 3C). Interestingly, rBUNV-N624Q titre was reduced in both A549 and C6/36 cells, but not to the same extent as rBUNV-Δ7 in C6/36 cells (Supplemental Fig 3C). Whilst these results show that glycosylation plays a role in viability of BUNV, they also show that glycosylation is not the reason for the lack of assembled rBUNV-Δ7 viruses in insect cells.

A final potential difference between rBUNV-WT and rBUNV-Δ7 was the lack of Gc head, which mediates trimerisation of the tripod. To determine whether spike tripod formation is important for BUNV assembly in insect cells, we identified potential head trimerisation residues in Gc (S575, R583, Q589 and D592) using PDBePISA (Proteins, Interfaces, Structures and Assemblies) and substituted each for alanine in the rBUNV-WT context to generate infectious mutant rBUNV-Δtripod. As above, we infected A549 and C6/36 cells with rBUNV-Δtripod at an MOI of 1 and harvested supernatant at 24 hpi (Supplemental Fig 3C). There was a slight reduction in rBUNV-Δtripod titre, in both cell lines, when compared to rBUNV-WT but again this was not as significant a reduction as rBUNV-Δ7 in C6/36 cells, suggesting lack of head domain trimerisation was not the reason that rBUNV-Δ7 viability is reduced in C6/36 cells.

### Generation of rBUNV-WT and rBUNV-Δ7 with HA-tagged Gc

Due to the only difference between rBUNV-WT and rBUNV-Δ7 being the 347 amino acid deletion in Gc, we next investigated whether this change affected Gc expression, processing, trafficking or localization, as these attributes could contribute to loss of virion production. To do this, we generated infectious rBUNV-WT and rBUNV-Δ7 variants expressing HA-tagged versions of their corresponding Gc proteins. The HA tag was introduced into the head region of WT-Gc and the remaining stalk region of Δ7-Gc (Supplemental Fig 1C) and successful rescue of corresponding viruses rBUNV-WT-Gc-HA and rBUNV-Δ7-Gc-HA was confirmed by detection of NP and Gc-HA by western blot analysis of both transfected and infected cell lysates (Supplemental Fig 1D).

The Gc-HA-tagged viruses were used to investigate expression of Gc in insect cells. Here, C6/36 cells were infected with either rBUNV-WT-Gc-HA or rBUNV-Δ7-Gc-HA at an MOI of 5 and lysate and supernatant samples were collected every 3 hpi until 24 hpi. Again, NP expression occurred slightly earlier in rBUNV-Δ7-Gc-HA-infected C6/36 cells when compared to rBUNV-WT-Gc-HA, but densitometry analysis showed this was not statistically significant (Fig 4A-B). Initial Gc-HA expression in rBUNV-Δ7-HA-infected C6/36 cells was seen at 12 hpi, similar to the initial expression of NP, and by 24 hpi there was robust expression of Gc-HA at the expected molecular weight (Fig 4C), suggesting that despite the deletion of the head/stalk domain, Gc-HA was correctly expressed and processed from the GPC precursor in C6/36 cells. Interestingly, Gc-HA expression in rBUNV-WT-Gc-HA-infected C6/36 cells was significantly slower than rBUNV-Δ7-Gc-HA, with initial expression first detected around 18 hpi, approximately 6 hours after rBUNV-Δ7-Gc-HA. However, by 21 and 24 hpi, there was no significant difference in the expression of WT or Δ7 Gc-HA. The supernatant samples showed that rBUNV-WT-Gc-HA was not affected by the addition of the HA tag, since titres are comparable to those of rBUNV-WT, and confirmed that the titres of rBUNV-Δ7-Gc-HA are similarly significantly reduced in C6/36 cells (Fig 4D).

**FIG 4.**
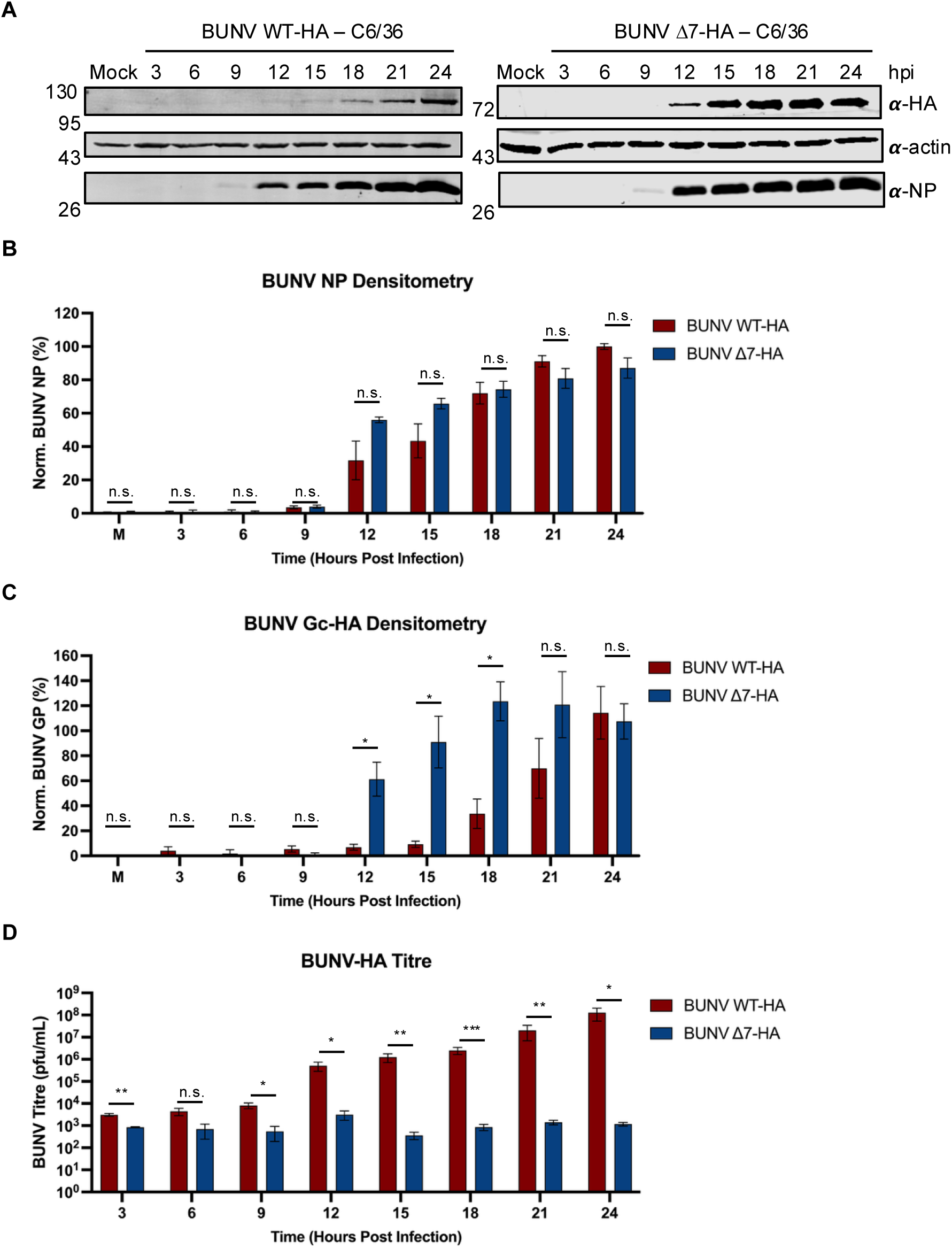
Comparison of growth kinetics and protein expression of wildtype rBUNV-HA and Δ7 rBUNV-HA in C6/36 cells. C6/36 cells were either mock-infected or infected with rBUNV-WT-Gc-HA (WT-HA) or mutant rBUNV-Δ7-Gc-HA (Δ7-HA) at an MOI of 5 and supernatants and cell lysates were collected at every 3 hours post infection (hpi) until 24 hpi (n=3). (A) Cell lysates were subject to western blot analysis and probed for expression of NP, HA (for expression of Gc-HA) and actin, as a loading control. (B) Densitometry analysis performed on (A) as expression of NP relative to actin. The expression of BUNV NP was normalised to rBUNV-WT-Gc-HA NP at 24 hours post infection. (C) Densitometry analysis performed on (A) as expression of Gc-HA relative to actin. The expression of BUNV Gc-HA was normalised to rBUNV-WT-Gc-HA Gc-HA at 24 hours post infection. (D) Titration of the supernatants was performed by plaque assay and titres were plotted on a logarithmic scale graph. All results were analysed by Student’s t-test (unpaired) whereby n.s. = p>0.05, * = p<0.05; ** = p<0.01; *** = p<0.001, comparing WT-HA to Δ7-HA at each timepoint.

### Δ7-Gc is transported to the plasma membrane in insect cells

To investigate whether Gc localisation is affected by deletion of the head/stalk region, C6/36 cells were infected with either rBUNV-WT-Gc-HA or rBUNV-Δ7-Gc-HA, fixed at 18 hpi and 24 hpi, stained for NP and Gc-HA and imaged using widefield microscopy (Fig 5). 18 hpi was chosen to reflect a time when WT-Gc expression was detectable but there was a significant difference between the levels of WT-Gc and Δ7-Gc expression, whereas 24 hpi was chosen to reflect a time when the difference between WT-Gc and Δ7-Gc abundance was not significant (Fig 4C). At 18 hpi and 24 hpi, NP distribution in rBUNV-WT-Gc-HA- and rBUNV-Δ7-Gc-HA-infected cells was indistinguishable, presenting as a diffuse cytoplasmic staining (Fig 5A-D, green).

**FIG 5.**
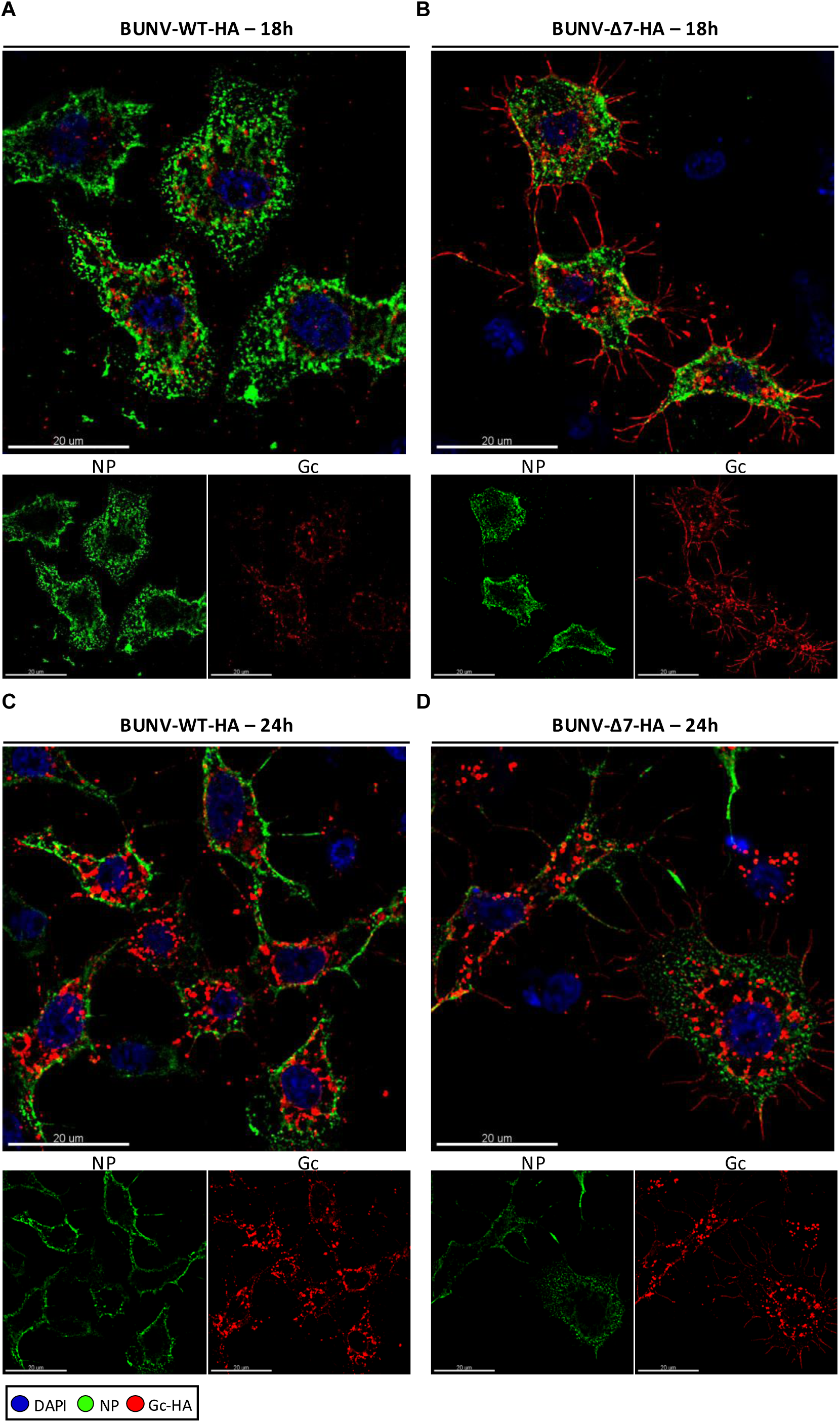
Widefield microscopy images of C6/36 cells infected with wildtype rBUNV-HA or Δ7 rBUNV-HA. C6/36 cells were infected with rBUNV-WT-Gc-HA (A and C) or rBUNV-Δ7-Gc-HA (B and D) at an MOI of 5 and fixed with formaldehyde at either 18 (A and B) or 24 (C and D) hpi. The cells were then permeabilized, blocked, and stained for the nucleus (DAPI, blue), BUNV NP (green), and BUNV Gc-HA (red) by indirect immunostaining. The cells were imaged on an Olympus IX83 widefield microscope at ×100 magnification. Z stack images were collected, and deconvolution (with 3 iterations) was performed. The slice showing the strongest signal was chosen for the display image. Scale bars representing 20 µm are shown. DAPI, 4’,6-diamidino-2-phenylindole.

However, in contrast to NP, localization of WT-Gc-HA and Δ7-Gc-HA in virus infected cells exhibited clear differences. At 18 hpi, WT-Gc-HA expressed in rBUNV-WT-Gc-HA-infected C6/36 cells appeared to localise exclusively within dense punctate regions, likely Golgi, in agreement with previous work (López-Montero & Risco, 2011) (Fig 5A-D, red). However, distribution of Δ7-Gc-HA at 18 hpi was different, with strong staining at the plasma membrane (Fig 5B, red) in addition to the aforementioned cytoplasmic puncta. At 24 hpi, when overall Gc expression levels are similar for WT-Gc-HA and Δ7-Gc-HA, staining of the respective WT-Gc-HA and Δ7-Gc-HA spikes was predominantly within cytoplasmic puncta (Fig 5C-D, red) however, Δ7-Gc-HA was still abundantly detected at the plasma membrane, with a much weaker signal for WT-Gc-HA.

To test whether Δ7-Gc was indeed present at the surface of the plasma membrane, C6/36 cells were infected with either rBUNV-WT-Gc-HA or rBUNV-Δ7-Gc-HA and fixed at 18 hpi and 24 hpi. Then, without permeabilization, the cells were stained for surface expression of Gc-HA and imaged using widefield microscopy (Fig 6). At both 18 hpi and 24 hpi timepoints, Δ7-Gc-HA showed a strong localisation to the plasma membrane (Fig 6B and D), whereas WT-Gc-HA showed a more punctate localisation at the plasma membrane, potentially representing released virions (Fig 6A and C). Together, these observations suggest the cellular distribution of Δ7-Gc-HA and WT-Gc-HA is different, potentially due to differences in protein processing, respective trafficking pathways, and/or pathway kinetics.

**FIG 6.**
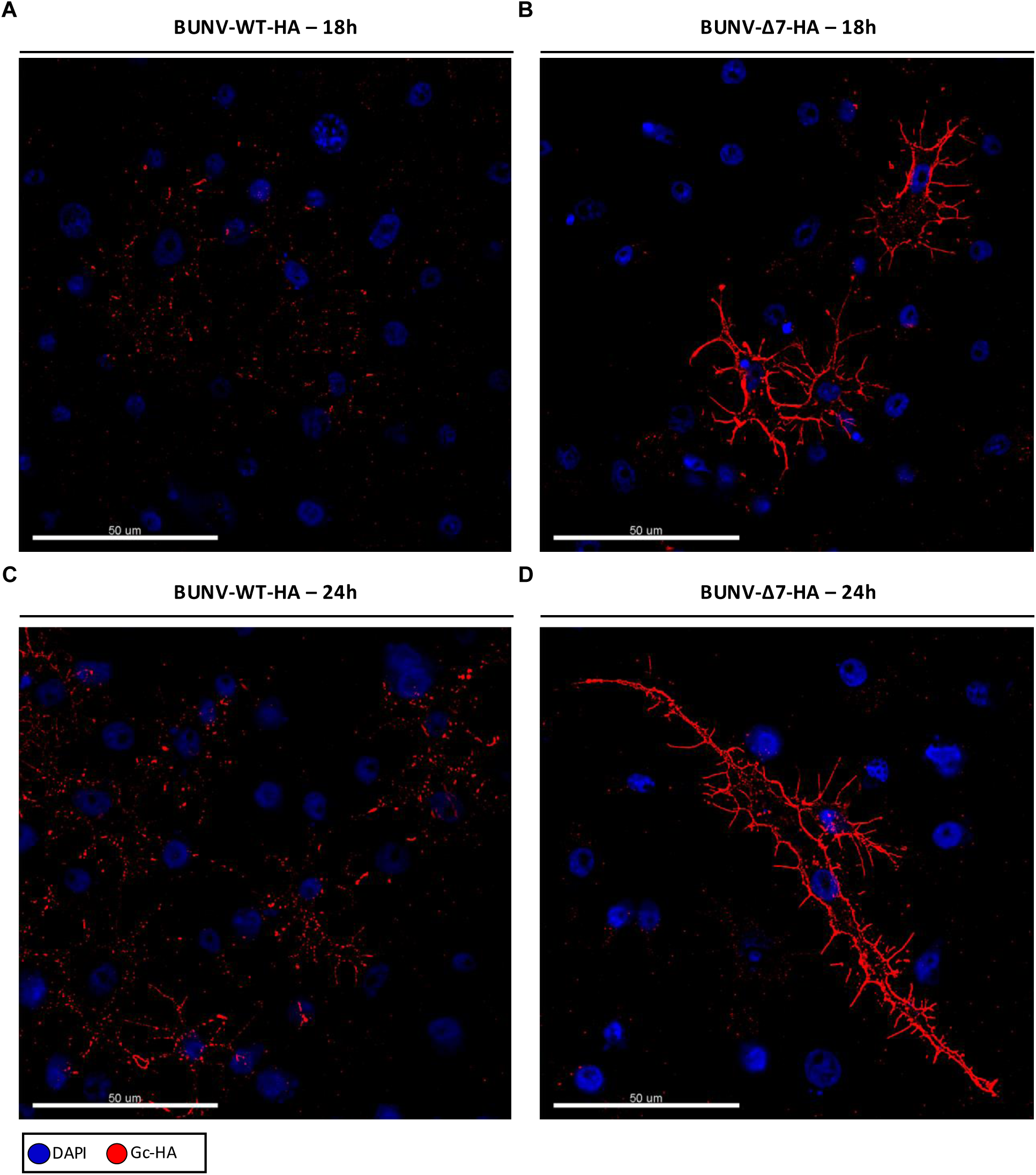
Widefield microscopy images of non-permeabilised C6/36 cells infected with wildtype rBUNV-HA or Δ7 rBUNV-HA. C6/36 cells were infected with rBUNV-WT-Gc-HA (A and C) or rBUNV-Δ7-Gc-HA (B and D) at an MOI of 5 and fixed with formaldehyde at either 18 (A and B) or 24 (C and D) hpi. The cells were not permeabilized, but were blocked, and stained for the nucleus (DAPI, blue) and BUNV Gc-HA (red) by indirect immunostaining. The cells were imaged on an Olympus IX83 widefield microscope at ×100 magnification. Z stack images were collected, and deconvolution (with 3 iterations) was performed. The slice showing the strongest signal was chosen for the display image. Scale bars representing 50 µm are shown. DAPI, 4’,6-diamidino-2-phenylindole.

### The head/stalk Gc deletion decreases Gc-NP association in insect cells, potentially blocking RNP recruitment

The results described above showed that Δ7-Gc-HA and WT-Gc-HA exhibit differences in their sub-cellular distributions within infected cells, and we hypothesized this could affect interactions with other virion components required for the assembly of infectious virions. To test this, we used co-immunoprecipitation followed by mass spectrometry analysis to identify intracellular binding partners of Δ7-Gc-HA and WT-Gc-HA. Briefly, A549 or C6/36 cells were infected with rBUNV-WT-Gc-HA or rBUNV-Δ7-Gc-HA at an MOI of 1 and incubated for 24 hpi, when cell lysates were collected. The lysates were then incubated with an HA antibody, following overnight incubation with Protein G-coupled magnetic beads. After washing, the beads were collected and subject to mass spectrometry analysis (Fig 7) or western blot analysis (Supplemental Fig 4). Initially, western blot (Supplemental Fig 4A) and densitometry analysis (Supplemental Fig 4B) were performed on the immunoprecipitated samples, staining for HA and NP expression. This suggested that Δ7-Gc-HA was less efficient than WT-Gc-HA at co-precipitating NP in C6/36 cells but not in A549 cells. However, densitometry analysis was not statistically significant, possibly due to differences in band intensity, smearing and molecular weight of WT-Gc-HA and Δ7-Gc-HA, plus the presence of contaminating antibody bands at the same molecular weight as NP.

**FIG 7.**
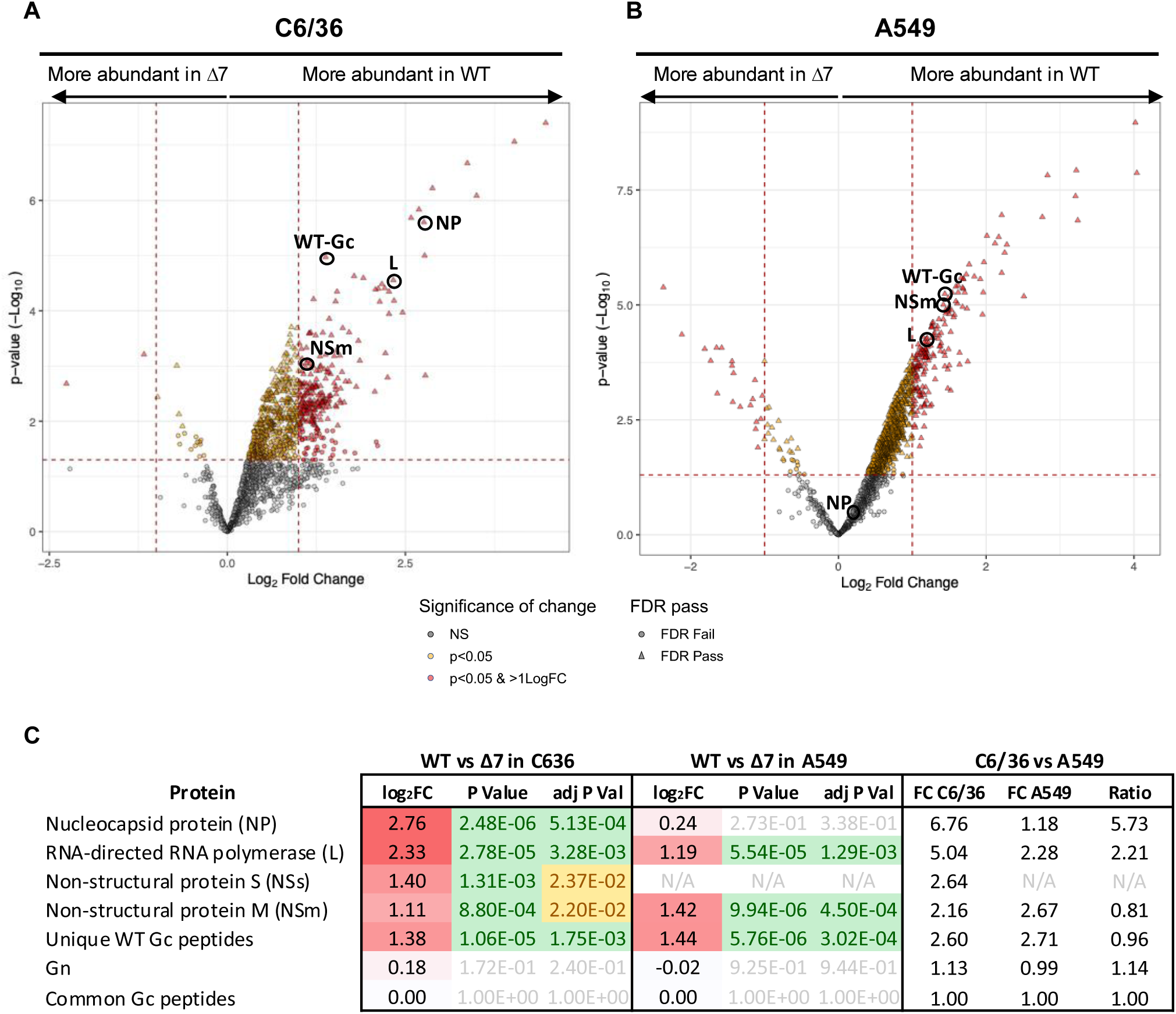
Mass spectrometry analysis of proteins identified from immunoprecipitation with BUNV Gc-HA. A549 cells or C6/36 cells were mock-infected or infected with rBUNV-WT-Gc-HA (WT-HA) or mutant rBUNV-Δ7-Gc-HA (Δ7-HA) at an MOI of 1 for 24 hours post infection. TMT-MS was then performed as described in methods, and normalised abundances were statistically analysed using the limma package. The volcano plots for the results from C6/36 cells (A) and A549 cells (B) were created by plotting the -log_10_ p-value of each protein against the log_2_ fold change. Proteins where p<0.05 are highlighted in yellow, and proteins where p<0.05 and log_2_FC is >1 or <-1 are highlighted in red. (C) Log_2_FC, p-value and adjusted p-value (adj p-val) for each viral protein identified in both cell lines is shown. Log_2_FC has been colour-coded as a gradient whereby 2=red and 0=white. The p-values and adj p-values have been colour-coded whereby green indicates p<0.01, yellow indicates p<0.05 and white indicates p>0.05. N/A means that no peptides were identified for that protein in that cell line.

Tandem mass-tag liquid chromatography mass spectrometry (TMT-MS) was performed on bead fractions precipitated from three biological repeats of either mock-, rBUNV-WT-Gc-HA- or rBUNV-Δ7-Gc-HA-infected A549 cells or C6/36 cells, allowing relative quantification of the interacting proteins of WT-Gc-HA and Δ7-Gc-HA. The log_2_- transformed abundances were normalised to the abundance of shared Gc sequences present in each condition (Fig 7C; Common Gc peptides), to compensate for any potential differences in the immunoprecipitation of WT-Gc-HA and Δ7-Gc-HA. Log_2_ fold changes (log_2_FC) were calculated for each protein, resulting in positive values (indicating WT-Gc-HA precipitated more protein than Δ7-Gc-HA), values close to 0 (indicating equal precipitation between WT-Gc-HA and Δ7-Gc-HA) or negative values (indicating Δ7-Gc-HA precipitated more protein than WT-Gc-HA) (Fig 7C). Volcano plots were then generated for each cell line to reflect log_2_FC changes and significance (p-value) (Fig 7A-B; Supplemental Dataset 1).

The analysis revealed a significantly higher abundance of NP peptides present in rBUNV-WT-Gc-HA-infected C6/36 cells compared to rBUNV-Δ7-Gc-HA-infected C6/36 cells (log_2_FC = 2.76, adj p-value = 5.13E-04). Interestingly, the log_2_FC of NP peptides present in rBUNV-WT-Gc-HA-compared to rBUNV-Δ7-Gc-HA-infected A549 cells was 0.24 (adj p-value = 3.38E-0.1), suggesting there was no significant difference between the precipitation of NP by WT-Gc-HA or Δ7-Gc-HA in A549 cells. Furthermore, this was not due to a lack of NP precipitation by Gc-HA because there was a similar significant enrichment of NP present in both WT and Δ7 A549 infections over mock (WT; log2FC = 3.93, p-value = 3.20E-03, Δ7; log2FC = 4.04, adj p-value = 4.83E-03) (Supplemental Dataset 1). There were also similar changes in the other bunyaviral proteins (L, NSm and NSs) in which the WT-Gc-HA was able to precipitate higher or similar abundances than Δ7-Gc-HA, to a greater extent in C6/36 cells than A549 cells (Fig 7C). In contrast to this, Gn had close to 0 log_2_FC values in both cell lines, suggesting Gn was equally precipitated by WT- and Δ7-Gc-HA. Finally, the Gc sequences unique to WT-Gc-HA were similarly enriched in both cell lines (C6/36; log_2_FC = 1.38, adj p-value = 1.75E-03, A549; log_2_FC = 1.44, adj p-value = 3.02E-04; WT Gc peptides in Fig 7C), which is to be expected due to the lack of some of these peptides in Δ7-Gc-HA. Together, this suggests that the ability of Δ7-Gc-HA to interact with NP is dramatically reduced in C6/36 cells, but not in A549 cells, consistent with the species-specific deficiency in insect host virion production.

## DISCUSSION

OBV glycoproteins are multifunctional, necessary for viral entry, assembly, exit and evasion of the host immune system. Remarkably, OBVs bearing large deletions within the Gc glycoprotein head and stalk domains can be isolated from infected animals, whereas these domains appear to be strictly conserved in OBV isolations from arthropods.

To better understand this discrepancy, we investigated the behaviour of a BUNV variant lacking the majority of the Gc head and stalk domains (rBUNV-Δ7; Shi et al., 2009) in cultured mammalian and insect cells, as well as live mosquitoes. We showed that rBUNV-Δ7 was capable of entering both mammalian and insect cells, and performing broadly equivalent gene expression to that of rBUNV-WT. However, whereas rBUNV-Δ7 generated equivalent mammalian cell titres to BUNV-WT, it was profoundly blocked in infectious virion production in both cultured C6/36 cells and *Aedes aegypti* mosquitoes. Thus, our results show the OBV Gc head and stalk domains play a species-specific role in infectious virion assembly.

To understand the molecular basis for why Δ7-Gc caused a species-specific block in infectious virion production, we tested host cell-based differences such as cholesterol content and optimal growth temperature, alongside viral Gc-specific differences including differential glycosylation potential and ability to form the characteristic OBV tripodal spike. While none of these variables influenced infectious virion production, we showed that deletion of the head/stalk influenced Gc cellular localisation, with increased abundance of Δ7-Gc at the plasma membrane, something which was not as evident with WT-Gc.

To investigate whether this differential localization of Δ7-Gc and WT-Gc was responsible for lack of rBUNV-Δ7 virion assembly in the insect host, we co-immunoprecipitated both Δ7-Gc and WT-Gc from insect cell lysates, revealing a significant reduction in the interaction between Δ7-Gc and RNP components NP and L, compared to WT-Gc. These reduced interactions are consistent with a role of Gc in RNP recruitment into newly generated virions in insect cells.

Crucially, immunoprecipitation of Δ7-Gc and WT-Gc from mammalian cell lysates did not result in differential pull-down of RNP components, a finding also consistent with the similar titres of released rBUNV-WT and rBUNV-Δ7 viruses in these cells. Taken together, these findings indicate that the presence of the Gc head and stalk leads to different infection outcomes in mammalian and insect cells.

One possible reason for the species-specific infection outcome could be the abundance of a cell factor specific to either insect or mammalian cells that facilitates the Gc-RNP interaction. Related to this, the TMT-MS analysis identified several insect proteins that had significantly reduced interactions with Δ7-Gc compared to WT-Gc. These included components of the COPI coatomer complex and various mitochondrial proteins, associated with vesicular transport or potential virus factory function, respectively. While these associations are provocative, their role in virion assembly remains to be determined. Interestingly, several other proteins with reduced Δ7-Gc interactions were categorised as uncharacterised in terms of function, with no mammalian homologues, which is suggestive of insect cell-specific pathways that could potentially play important roles in virion assembly (Supplemental Dataset 1). Further investigation could offer insight into how viral assembly is directed in insect cells.

Alternatively, the Gc head and stalk could dictate species-specific infection outcomes through differences in Gc expression kinetics or trafficking, where loss of the head/stalk could affect the availability of spike proteins to interact with RNPs at destined assembly sites. In agreement with this, we showed Δ7-Gc synthesis occurred sooner than for WT-Gc in insect cells, and exhibited an altered cellular distribution, showing increased abundance at the plasma membrane, as well as cytoplasmic puncta. It could be that deletion of the head/stalk may cause the loss of an uncharacterised insect cell-specific signal that has altered Gc retention, processing or maturation.

A similar scenario was proposed for Crimean Congo haemorrhagic fever virus (CCHFV), for which deletion of GP38 affected Gc maturation and reduced Gc localisation to the Golgi, and deletion of NSm increased the rate of Gc trafficking and resulted in reduced N-glycosylation (Freitas et al., 2020). Similarly, the Gc head/stalk perhaps play a role in directing its own maturation and/or trafficking, both of which could result in a reduced interaction with NP and affect assembly.

It is interesting to speculate how removal of the Gc head might affect interaction with NP for virion assembly, particularly as the current model describing the BUNV assembly site places the Gc head and tail within the Golgi lumen, and NP within the cytosol. This being the case, it is highly unlikely that the Δ7-Gc deletion prevents a direct interaction between NP and its head/stalk. The Gc-NP interaction is thought to be driven by the cytoplasmic tails of Gn and Gc, fulfilling the role of a canonical viral matrix protein and mediating recruitment of all viral components necessary for virion generation (Shi et al., 2007). Our TMT-MS analysis showed that the interaction between Δ7-Gc and Gn was not significantly different to that of WT-Gc, which confirms removal of the Gc head does not affect the heterodimeric complex. However, whether trimerisation of the floor domain, or the Gc and Gn cytoplasmic tails, is disrupted is not known. In an attempt to decipher this, we generated a rBUNV-Δtripod mutant to mimic the lack of trimerisation driven by the Gc head. This produced high titres from C6/36 cells, unlike rBUNV-Δ7, suggesting lack of tripod formation (i.e. trimerisation of the head domain) is not required for virus assembly. There is also the potential that the complex glycosylation seen in mammalian cells is required to direct proper folding and multimerization of Gc, even in its truncated version, whereas the simpler glycosylation in insect cells might not be sufficient to drive correct folding. However, Δ7-Gc is missing only one glycosylation site compared to WT-Gc, and mutant virus rBUNV-N624Q showed that removal of this site did not result in the same low titres as rBUNV-Δ7 from insect cells. The deleted sequence corresponds to nearly 350 amino acids and therefore it is possible that there are other unknown modification sites or signals, potentially insect-cell-specific, present in this region.

The observation that the Gc head and stalk domains are dispensable for entry in both mammalian and insect cells suggests that either removal of the Gc head does not disrupt receptor binding/attachment, or that exposure of the fusion domain offers a receptor-independent entry route, at least *in vitro*. The redundancy of these domains in mammalian cells is consistent with the low sequence conservation of the OBV Gc N-terminus (Coupeau et al., 2013; Fischer et al., 2013; Iroegbu & Pringle, 1981; Pollitt et al., 2006; Shi et al., 2009; Varela et al., 2016), with this antigenic drift likely driven by the need to escape neutralizing antibodies that target these regions (Hellert et al., 2019; Stubbs et al., 2021; Thannickal et al., 2023) Taken together, the role of the OBV Gc head and stalk regions are likely to offer a prominent target to the mammalian adaptive immune response, misdirecting antibody-mediated neutralization from the functionally-critical residues involved in membrane fusion that are located in the Gc C-terminus, and whose sequence is more rigid due to a high functional constraint (Roman-Sosa et al., 2016). However, the results presented here suggest the Gc head and stalk also provide further critical roles in the virus multiplication cycle aside from a neutralizing antibody decoy.

Our results offer a plausible explanation for why naturally occurring SBV mutants bearing deletions of the Gc head domain can be detected in the ruminant foetus, but not in infected viraemic mothers, nor in insects (Coupeau et al., 2013; Fischer et al., 2013; Wernike et al., 2021; Wernike & Beer, 2019). We propose that Gc head/stalk deletion mutants arise due to an inherent mutational hotspot in the Gc N-terminus, sometimes leading to genetic drift, but also sometimes leading to large scale sequence deletion of the entire head, often along with deletions in the stalk. In immune competent ruminant mothers, such deletions would likely be selected against due to exposure of functionally critical C-terminal Gc sites, and subsequent antibody-mediated neutralization. In contrast, due to the lack of a robust adaptive immune response, such head/stalk deleted mutants are capable of rapid amplification in the early-stage foetus, resulting in the devastating pathogenicity associated with neonatal OBV infection. In this context, while the strong selection pressure for maintenance of the Gc head/stalk in the infected ruminant may be a contributing factor, we propose that the lack of detection of head/stalk deleted OBVs in insects primarily results from the species-specific assembly deficiency described here. Interestingly, Gc may not be the only OBV protein able to cause a species-specific block in virus multiplication. A recent report describes an SBV mutant bearing a single NP mutation as responsible for a lack of virus multiplication within both cultured *Culicoides* midge cells (Sick et al., 2024) as well as live midges (Wernike et al., 2025). Taken together, these findings highlight major knowledge gaps exist in understanding the complex interplay of determinants of virus fitness in the dual host OBV multiplication cycle.

## MATERIALS AND METHODS

### Cell lines

BHK-21 cells, BSR-T7 cells and A549 cells were acquired from ATCC and maintained in high-glucose Dulbecco’s modified Eagle medium (DMEM; Sigma-Aldrich), supplemented with 10 % heat-inactivated foetal bovine serum (FBS), 100 µg of streptomycin/mL and 100 U of penicillin/mL, and incubated in a humidified incubator at 37°C with 5 % CO_2_. BSR-T7 cells constitutively expressing T7 RNA polymerase (T7P) (Buchholz et al., 1999) were also additionally supplemented with G418 (500 µg/mL) every other passage to maintain the T7P plasmid. C6/36 insect cells were gifted by Andrew Tuplin (University of Leeds) and were maintained at 28°C in Leibovitz’s L-15 media (Thermo Fisher Scientific) which was supplemented with 10 % Tryptose Phosphate Broth (Thermo Fisher Scientific) and 100 µg of streptomycin/mL and 100 U of penicillin/mL.

### Plasmids

Plasmids encoding the full-length cDNAs that represent the intact S (pT7riboBUNS(+); Genbank accession number: NC_001927.1), M (pT7riboBUNM(+); Genbank accession number: NC_001926.1) and L (pT7riboBUNL(+); Genbank accession number: NC_001925.1) segments of BUNV were gifted by Alain Kohl (University of Glasgow) (Bridgen & Elliott, 1996). Site-directed mutagenesis was used to delete amino acids D480 to N826 from pT7riboBUNM(+) to generate pT7riboBUNMΔ7(+), as previously described (Shi et al, 2009).

Plasmids pT7riboBUNM-HA(+) and pT7riboBUNMΔ7-HA(+) were synthesised through PCR insertion of the HA tag (YPYDVPDYA), flanked by flexible 4xG/S sequences, into the Gc region (inserted between amino acids T502 and D503 in pT7riboBUNM-HA(+) or between amino acids V482 and Y483 in pT7riboBUNMΔ7-HA(+) (equivalent residues in WT are V829 and Y830). Plasmid pT7riboBUNM-trimer(+) was synthesised through PCR alanine substitution of residues S575, R583, Q589, D592. Plasmid pT7riboBUNM-N624Q (+) was synthesised by PCR substitution of residue N624 to glutamine. All primers sequences are available upon request.

### Generation of recombinant BUNV from cDNA

BSR-T7 cells were seeded in 6-well plates at a density of 2×10^5^ cells/well. Following overnight incubation, the cells were transfected with 1 µg of pT7riboBUNS(+), 1 µg of pT7riboBUNM(+), 1 µg of pT7riboBUNL(+) and 0.3 µg of pUC57-T7 in 200 µL OptiMEM media, followed by 2.5 µL/µg TransIT-LT1 transfection reagent (Mirus). For recovery of the Δ7 variant, pT7riboBUNM(+) was replaced with pT7riboBUNMΔ7(+). Tagged or mutant versions of viruses involved replacing the appropriate plasmids by their tagged or mutated counterparts. Control transfections were also performed, in which pT7riboBUNL(+) was excluded. At 4-hour post transfection (hpt), the media was removed and replaced with 2 mL of DMEM supplemented with 2.5 % FBS. At 120 hpt, the supernatants were collected, clarified and used to infect BHK-21 cells, seeded at a cell density of 5×10^5^ cells in T25 flasks the previous day. 48-72 hours post infection (hpi), upon evidence of cytopathic effect, supernatant was collected, centrifuged at 4000 *x g* for 15 mins, filtered through a 0.45 µm filter and aliquoted into cryo-vials for storage at -80 °C. To prepare bulk stocks of virus, T175 flasks were seeded with 1.5×10^7^ BHK-21 cells, incubated for 24 h and then infected with BUNV (or the BUNV variants) at an MOI of 0.01. At 48 hpi, supernatant was clarified and filtered as described above.

### Viral protein expression and virus release in mammalian and insect cells

A549 (1×10^5^ cells/well), BHK-21 (1×10^5^ cells/well) and C6/36 (4×10^5^ cells/well) were seeded in 12-well plates and incubated at either 37 °C (mammalian) or 28 °C (insect). The cells were then infected with rBUNV-WT or rBUNV-Δ7 at an MOI of 5 and incubated for a total of 24 hpi. Supernatant and cell lysate samples were collected at 6 hpi, 12 hpi and 24 hpi for determination of virus release and NP expression respectively. For more detailed analysis, C6/36 (4×10^5^ cells/well) were seeded in 12-well plates and incubated at 28 °C. The cells were then infected with rBUNV-WT-Gc-HA or rBUNV-Δ7-Gc-HA at an MOI of 5 and incubated for a total of 24 hpi. Supernatant and lysate samples were collected every 3 hours following infection for determination of virus release and NP and Gc-HA expression respectively.

For collection of intracellular and extracellular virus samples, A549 (1.5×10^5^ cells/well) and C6/36 (4×10^5^ cells/well) were seeded in 12-well plates, infected with rBUNV-WT or rBUNV-Δ7 at an MOI of 5 and incubated for 24 hours at either 37 °C (mammalian) or 28 °C (insect). Supernatants (extracellular – 1 mL) were collected into cryo-vials and stored at -80 °C. Cells (intracellular) were washed twice with 1X TNE (10 mM Tris-HCl, pH 7.4, 100 mM NaCl, 1 mM EDTA) and then scraped into 1 mL 1X TNE and freeze-thawed three times to lyse the cellular membranes. Both extracellular and intracellular samples were titred by crystal violet plaque assay.

To examine BUNV behaviour at lower incubation temperatures, A549 (2×10^5^ cells/well) or BHK-21 (2×10^5^ cells/well) were seeded in 12-well plates and incubated at 30 °C. The cells were then infected with rBUNV-WT or rBUNV-Δ7 at an MOI of 5 and incubated for a total of 24 hpi. Supernatant samples were collected at 24 hpi for determination of virus titre (*n=1*).

To determine the effect of mutating the trimerisation residues or the glycosylation site missing in rBUNV-Δ7, A549 (1.5×10^5^ cells/well) and C6/36 (4×10^5^ cells/well) were seeded in 12-well plates, infected with either rBUNV-WT, rBUNV-Δ7, rBUNV-Δtripod or rBUNV-N624Q at an MOI of 1 and incubated for 24 hours at either 37 °C (mammalian) or 28 °C (insect). Supernatant samples were collected at 24 hpi for determination of virus titre (*n=2*).

### Mosquito infection

Female *Aedes aegypti* mosquitoes (Poza Rica, Mexico) were sorted and housed in cardboard pint cups the day prior to infection. Following a 24-hour starvation period, the mosquitoes were fed a blood meal of washed sheep’s blood containing either rBUNV-WT or rBUNV-Δ7 at 1 x 10^7^ PFU/mL, supplemented with 5 mM ATP. Engorged mosquitoes were transferred to new cups with 10 % sucrose and kept at 28 °C and 70 % humidity for seven days, at which point legs and wings were harvested in microcentrifuge tubes containing glass beads and 250 μL DMEM supplemented with 2 % newborn calf serum (NBCS). Bodies were also collected in 250 μL DMEM with 2 % NBCS. Samples were ground using a Pro Series Bullet Blender homogenizer for one minute at maximum speed and then centrifuged at 10,000 *x g* for three minutes. Plaque assays were conducted on all samples to determine viral titres, as follows. Supernatant from each homogenized mosquito sample was serially diluted 10-fold in DMEM and inoculated onto a monolayer of Vero cells. Cells were incubated at 37 °C and 5 % CO_2_ for one hour prior to being overlaid with 1.5 mL DMEM supplemented with 0.8 % agarose, 2 % NBCS, and 1 % Antibiotic-Antimycotic. Cells were incubated at 37 °C for three days and then fixed with 4 % formalin for one hour before the overlay was removed and cells were stained with crystal violet to allow plaques to be counted. Viral titres were quantified by counting plaques on the lowest countable dilution.

Day 0 samples were collected from entire mosquitoes harvested immediately after feeding on the infectious blood meal. Samples were homogenized and clarified as detailed above. Plaque assays were conducted to determine viral titers.

For protein analysis, entire mosquitoes were harvested in 2 X Laemmli buffer 7 days post-infection and ground using a pestle for western blotting.

### Cholesterol repletion in insect cells

To understand whether the lack of cholesterol in insect cells was affecting rBUNV-Δ7 assembly, C6/36 (4×10^5^ cells/well) were seeded on glass coverslips in 12-well plates and incubated for 24 hours at 28 °C. Prior to infection, the cells were pre-treated with either methyl-β-cyclodextrin (MBCD; 85 µM; vehicle control), or 0.05 mg/mL or 0.1 mg/mL cholesterol, for 1 hour. The cells were then washed three times with 1X PBS. Next, rBUNV-WT or rBUNV-Δ7 at an MOI of 5 in serum-free Leibovitz’s L-15 media was added to the cells and incubated at 28 °C for 1 hour. The cells were then washed three times with 1X PBS and incubated for 24 hours at 28 °C in serum-free Leibovitz’s L-15 media. Supernatant samples were collected at 24 hpi for determination of virus titre (*n=1*).

### Virus purification

12 T175 flasks were seeded with 2.75×10^7^ C6/36 cells at least 24 hours before infection. Cells were washed once with 1x PBS and infected with rBUNV-Δ7, at a MOI of 0.01, for 1 hour in 3 mL serum free media at 37 °C (or 28 °C) before adding 15 mL 2.5 % DMEM (or complete Leibovitz’s L-15 media). At 48 hpi, supernatant was collected and centrifuged for 30 minutes at 4000 x g at 4 °C. The supernatant was filtered through a 0.45 µm filter and centrifuged again for 30 minutes at 4000 x g at 4 °C. The supernatant was then transferred to SW32 ultra-centrifuge tubes (Becker-Coltman) and 8 mL 20 % sucrose (supplemented with 1 X cOmpleteTM, Mini, EDTA-free Protease I inhibitor cocktail (Sigma-Aldrich)) was underlaid. Samples were centrifuged for 2 hours at 90,000 x g at 4 °C. The sucrose cushion and supernatant were removed after centrifugation and the virus pellet was air-dried for 5 minutes, upside down on tissue. 15 µL resuspension buffer (0.1 X PBS + 1 X cOmpleteTM, Mini, EDTA-free Protease I inhibitor cocktail (Sigma-Aldrich)) was added to the virus pellet and the ultra-centrifuge tubes were covered with two layers of parafilm, before incubating on a rocker overnight at 4 °C. The resuspended pellet was aliquoted and flash-frozen using liquid nitrogen and then stored at -80 °C. Samples were taken for silver stain analysis.

### Electron microscopy of insect cell slices

T25 flasks were seeded with 4×10^6^ C6/36 cells 24 hours prior to infection. Cells were washed once with PBS and either mock-infected or infected with rBUNV-WT or rBUNV-Δ7 at an MOI of 5 to ensure every cell was infected. The cells were incubated for 1 h at 28 °C, after which the infection media was replace with complete Leibovitz’s L-15 media and incubated at 28 °C for 24 hours. Cells were then fixed with 2.5 % glutaraldehyde (made in 1X PBS) and incubated for 2 h at room temperature. The cells were scraped into the media and then centrifuged at 300 x g for 5 mins to collect the cell pellet. The pellet was washed twice with 1X PBS and then treated with 1 % osmium tetroxide (made in 1X PBS) for 1 h depending on size and thickness of the sample. The pellet was washed twice with 1X PBS and then dehydrated with an ascending alcohol series (20 %, 40 %, 60 %, 80 %, 2X 100 %) for 20-60 min per change, depending on specimen size. The cells were then washed twice with propylene oxide for 20 min each and then treated with Araldite-propylene oxide (1:1) for several hours or overnight, Araldite-propylene oxide (3:1) for several hours and finally neat Araldite for 3-8 h (Luft, 1961). The cells were transferred to embedding moulds with fresh Araldite and left to polymerise overnight at 60 °C. In each treatment or wash, the sample was handled in small Eppendorf tubes and centrifuged (300 x g, 5 min) to change the solution and resuspended using a whirly mixer. Ultra-thin sections were prepared using a Reichert-Jung ultracut E ultramicrotome and picked up on 3.05 mm grids and then stained with saturated uranyl acetate (5 min – 120 min) and Reynolds lead citrate (5 min – 30 min) (Reynolds, 1963).

### Virus titration

Plaque assays were performed on confluent monolayers of BHK-21 cells. Serial dilutions of BUNV (10-fold) made in serum-free DMEM were added to the cells and incubated for 1 h at 37 °C. The dilutions were removed from the cells and overlaid with 1.2 % Avicel® (RC-581, FMC Biopolymer). After 3 days incubation at 37 °C, the cells were fixed through the overlay with 8 % formaldehyde in water for 10 mins. The fixative was washed off with water and then the cells were stained with 2 % crystal violet in 20 % ethanol. Plaques were counted and the titre was determined.

### Western blot analysis

Cells were lysed with 1X radioimmunoprecipitation assay (RIPA) buffer (50 mM Tris-HCl pH 7.5, 150 mM NaCl, 1 % (v/v) NP40 alternative, 0.5 % (w/v) sodium deoxycholate and 0.1 % sodium dodecyl sulphate (SDS; w/v)), supplemented with 1X Halt™ Protease Inhibitor Cocktail (100X; Thermo Fisher Scientific) for 15 min on ice. Lysates were collected; proteins were resolved on a 12 % or 15 % (immunoprecipitation samples) SDS-PAGE gel and then transferred to a polyvinylidene fluoride (PVDF) membrane. The transfer was performed at 15 V for 30 min, using the Trans-Blot turbo (Bio-Rad). After transfer, the membrane was blocked for 1 h in Odyssey blocking buffer (PBS) (Licor; diluted 1:1 with 1X PBS containing 0.1 % Tween^TM^ 20 [PBS-T]). Subsequently, the membrane was stained with the primary antibodies (made in 1:4 [blocking buffer: PBS-T]; anti-BUNV NP 1:5000, anti-actin 1:5000, anti-HA 1:5000) for 1 h rocking, at room temperature and then with corresponding secondary antibodies (made in 1:4 [blocking buffer: PBS-T]; all secondary antibodies used at 1:10000) for 1 h at room temperature. The membrane was washed and visualised on the Odyssey M imaging system (Licor). Densitometry analysis was performed using ImageJ over three independent experiments.

### Silver stain analysis

Viral pellet samples (10 µL) were resolved by 15 % SDS-PAGE and then fixed and stained using the ProteoSilver^TM^ Kit (Sigma-Aldrich), following the manufacturer’s protocol.

### Immunofluorescence analysis

To assess localisation of BUNV proteins during the virus replication cycle, C6/36 cells (4×10^5^) were grown on glass coverslips and infected with rBUNV-WT-Gc-HA (MOI = 5) or rBUNV-Δ7-Gc-HA (MOI = 5). At specified time points, the cells were fixed with 4 % formaldehyde for 10 min, permeabilised with 0.3 % triton-x-100 for 10 min and blocked with 1 % bovine serum albumin (BSA) for 1 h. Cells were then labelled with anti-BUNV NP (1:5000) and anti-HA (for Gc staining; 1:5000) for 1 h, followed by labelling with corresponding Alexa Fluor 488 nm or 647 nm secondary antibodies (all secondary antibodies used at 1:5000), respectively, for 1 h. Cells were then washed and mounted onto microscope slides using prolong gold containing DAPI (Invitrogen-Molecular Probes). Stained cells were viewed on the Olympus IX83 widefield microscope at 100 x magnification. Z stack images were collected, and deconvolution (with 3 iterations) was performed (CellSens, Olympus). The slice showing the strongest signal was chosen for the display image. For non-permeabilised samples, all steps were performed as above, with the permeabilization step omitted.

### Immunoprecipitation of Gc-HA and interacting proteins

A549 or C6/36 cells (1×10^7^ cells/flask) were seeded and incubated at either 37°C (mammalian) or 28 °C (insect). The cells were then infected with rBUNV-WT-Gc-HA or rBUNV-Δ7-Gc-HA at an MOI of 1 and incubated for a total of 24 hpi. Cells were washed once with 1X PBS and lysed in 1 mL immunoprecipitation lysis buffer (25 mM Tris-HCl, 150 mM NaCl, 1 mM EDTA, 1 % NP-40, 5 % glycerol), supplemented with 1X Halt™ Protease Inhibitor Cocktail (100X; Thermo Fisher Scientific). The cells were lysed for 30 min at 4 °C, after which the lysates were collected and centrifuged at 13,000 *x g* for 15 min. The lysates were incubated rotating overnight at 4 °C with Dynabeads™ Protein G beads, which had previously been incubated with anti-HA antibody (1:80) in PBS-T (0.02 % Tween^TM^ 20) for 2 h at room temperature and then washed with PBS-T to remove any unbound antibody. After overnight incubation, the beads were drawn out of solution using the DynaMag^TM^ magnetic rack (Invitrogen), and unbound protein was removed. The beads were subsequently washed three times with PBS-T and then resuspended in 50 µL PBS-T for mass spectrometry analysis.

### TMT labelling and high pH reversed phase (RP) chromatography

Mammalian and insect samples were analysed using separate TMT experiments, as described below. Immuno-isolated samples were reduced (10 mM TCEP, 55 °C for 1 h), alkylated (18.75 mM iodoacetamide, room temperature for 30 min) and then digested from the beads with trypsin (1.25 µg trypsin; 37 °C, overnight). The resulting peptides were then labelled with TMTpro sixteen-plex reagents according to the manufacturer’s protocol (Thermo Fisher Scientific, Loughborough, LE11 5RG, UK) and the labelled samples pooled and desalted using a SepPak cartridge according to the manufacturer’s instructions (Waters, Milford, Massachusetts, USA). Eluate from the SepPak cartridge was evaporated to dryness and resuspended in buffer A (20 mM ammonium hydroxide, pH 10) prior to fractionation by high pH reversed-phase chromatography using an Ultimate 3000 liquid chromatography system (Thermo Scientific). Samples were loaded onto an XBridge BEH C18 Column (130Å, 3.5 µm, 2.1 mm X 150 mm, Waters, UK) in buffer A and peptides eluted with an increasing gradient of buffer B (20 mM Ammonium Hydroxide in acetonitrile, pH 10) from 0-95 % over 60 minutes. The resulting fractions (concatenated into 4 in total) were evaporated to dryness and resuspended in 1 % formic acid prior to analysis by nano-LC MSMS using an Orbitrap Fusion Tribrid mass spectrometer (Thermo Scientific).

### Nano-LC mass spectrometry

High pH RP fractions were further fractionated using an Ultimate 3000 nano-LC system in line with an Orbitrap Fusion Tribrid mass spectrometer (Thermo Scientific). In brief, peptides in 1 % (vol/vol) formic acid were injected onto an Acclaim PepMap C18 nano-trap column (Thermo Scientific). After washing with 0.5 % (vol/vol) acetonitrile 0.1 % (vol/vol) formic acid, peptides were resolved on a 250 mm × 75 μm Acclaim PepMap C18 reverse phase analytical column (Thermo Scientific) over a 150 min organic gradient with a flow rate of 300 nl min−1. Solvent A was 0.1 % formic acid, and Solvent B was aqueous 80 % acetonitrile in 0.1 % formic acid. Peptides were ionized by nano-electrospray ionization at 2.0kV using a stainless-steel emitter with an internal diameter of 30 μm (Thermo Scientific) and a capillary temperature of 275 °C.

All spectra were acquired using an Orbitrap Fusion Tribrid mass spectrometer controlled by Xcalibur 2.1 software (Thermo Scientific) and operated in data-dependent acquisition mode using an SPS-MS3 workflow. FTMS1 spectra were collected at a resolution of 120,000, with an automatic gain control (AGC) target of 200,000 and a max injection time of 50 ms. Precursors were filtered with an intensity threshold of 5,000, according to charge state (to include charge states 2-7) and with monoisotopic peak determination set to peptide. Previously interrogated precursors were excluded using a dynamic window (60s +/-10 ppm). The MS2 precursors were isolated with a quadrupole isolation window of 1.2 m/z. ITMS2 spectra were collected with an AGC target of 10,000, max injection time of 70ms and CID collision energy of 35 %.

For FTMS3 analysis, the Orbitrap was operated at 50,000 resolution with an AGC target of 50,000 and a max injection time of 105 ms. Precursors were fragmented by high energy collision dissociation (HCD) at a normalised collision energy of 60 % to ensure maximal TMT reporter ion yield. Synchronous Precursor Selection (SPS) was enabled to include up to 10 MS2 fragment ions in the FTMS3 scan.

### Mass spectrometry data analysis

Raw data files were processed and quantified using Proteome Discoverer software v2.4 (Thermo Scientific) and searched against the UniProt Human database (downloaded January 2025: 83095 entries) or the UniProt *Aedes albopictus* (downloaded January 2025: 26006 entries) using the SEQUEST HT algorithm. For each experiment, the UniProt Bunyamwera virus database (downloaded January 2025: 7 entries) was also included in the searches. Peptide precursor mass tolerance was set at 10 ppm, and MS/MS tolerance was set at 0.6 Da. Search criteria included oxidation of methionine (+15.995 Da), acetylation of the protein N-terminus (+42.011 Da), methionine loss from the protein N-terminus (-131.04 Da) and methionine loss plus acetylation of the protein N-terminus (-89.03 Da) as variable modifications and carbamidomethylation of cysteine (+57.021 Da) and the addition of the TMT mass tag (+304.207 Da) to peptide N-termini and lysine as fixed modifications. Searches were performed with full tryptic digestion and a maximum of 2 missed cleavages were allowed. The reverse database search option was enabled, and all data was filtered to satisfy false discovery rate (FDR) of 5 %.

### Mass spectrometry statistical analysis

The MS data were searched against the human, *Aedes albopictus* and Bunyamwera virus databases retrieved 2025-01-24 and updated with additional annotation information on 2025-05-14. Protein groupings were determined by PD2.4, however, further the master protein selection was improved, and all following statistical analyses were performed in the R statistical environment version 4.4.0. The master protein improvement script first searches UniProt for the current status of all protein accessions and updates redirected or obsolete accessions. The script further takes the candidate master proteins for each group and uses current UniProt review and annotation status to select the best annotated protein as master protein without loss of identification or quantification quality.

The protein abundances for each sample were normalised using sum of peptide abundance before being Log_2_ transformed to bring them closer to a normal distribution. The data were statistically analysed using normalised abundances. Further analysis was then performed on data normalised to the proportion of common Gc peptides between the WT and Δ7 samples, to account for differences in Gc concentration between the two samples.

A single limma model was constructed for all conditions and moderated t-tests performed for each comparison of interest, then the p-value was adjusted using the Benjamini-Hochberg FDR method. Additionally, univariate unpaired t-tests were performed individually for each protein and each comparison of interest, followed by p-value adjustment using the Benjamini-Hochberg FDR method. The limma package (Ahlmann-Eltze & Anders, 2019) was used for statistical test as this has been recently identified as one of the highest scoring methods for identification of differentially expressed proteins (Peng et al., 2024).

### Mass spectrometry PCA

PCAs were calculated using the prcomp package, and the plotted using the ggplot2 package. Principal Components 1 and 2 were plotted against each other to give an indication of the main sources of variance, and 3 and 4 were plotted to infer any further trends for both raw and normalised abundances.

### Mass spectrometry volcano plots

For each comparison, using both raw and normalised data, the – Log_10_ p-value of each protein was plotted against the log2 fold change. Proteins where p<0.05 were highlighted in yellow, and proteins where p<0.05 and Log_2_FC>1, or where p<0.05 and Log_2_FC<-1 were highlighted in red.

### Statistical analysis

The statistical significance of data was determined by performing a Student’s *t* test. The statistical significance of the titre data from mosquitoes was determined by performing a Mann Whitney test. Significance was deemed when the values were less than or equal to the 0.05 *p* value.

## ACKNOWLEDGEMENTS

We acknowledge funding from MRC grant MR/T016159/1 (to JNB and JF) funding ABS and HNT, BBSRC grant BB/V007467/1 funding HP, UKHSA PhD studentship grant OB1 to OB. JF was supported by grant PID2023-149259NB-I00, funded by MICIU/AEI/ 10.13039/501100011033 and by “ERDF A way of making Europe”. The authors would like to thank Xiaohong Shi for extremely useful discussions about the results. The authors would also like to extend their thanks to Dr Kate Heesom and Dr Philip Lewis at the University of Bristol Proteomics facility for performing the TMT-MS experiments and subsequent analysis; to Dr Ruth Hughes and Dr Sally Boxall of the bioimaging facility, Faculty of Biological Sciences, University of Leeds for their expert assistance and use of the Olympus IX83; and to Martin Fuller of the Astbury Biostructure Laboratory electron microscopy facility, University of Leeds for his help preparing and imaging insect cell samples.

